# Inhibition of microtubule polymerization and dynein impairs the nuclear localization of the ependymoma-associated ZFTA-RELA fusion protein and NF-κB activation

**DOI:** 10.1101/2024.11.26.625308

**Authors:** Masaki Ishii, Naoki Suto, Kazuaki Katakawa, Kosho Makino, Sachiko Toma-Fukai, Shunsuke Sueki, Masahiro Anada, Hirotatsu Kojima, Shinya Ohata

## Abstract

Ependymomas are rare and chemotherapy-resistant gliomas. One subclass of supratentorial ependymomas, ST-EPN-ZFTA, expresses a fusion protein consisting of a nuclear protein, zinc finger translocation associated (ZFTA), and *v-rel* reticuloendotheliosis viral oncogene homolog A (RELA), an effector transcription factor of the nuclear factor-κB (NF-κB) pathway (ZFTA-RELA). Constitutive localization of ZFTA-RELA to the nucleus hyperactivates the oncogenic NF-κB signaling pathway, thereby contributing to the pathogenesis of ST-EPN-ZFTA. To identify compounds that inhibit NF-κB activity induced by ZFTA-RELA, we established a high-throughput screening system using the NF-κB-responsive luciferase reporter cell line 6E8, which expresses ZFTA-RELA in a doxycycline-dependent manner. A chemical library of 9600 compounds selected for their structural diversity was screened, and a colchicine derivative was identified. Among colchicine and six of its derivatives, the IC_50_ on ZFTA-RELA-dependent NF-κB-responsive luciferase activity in 6E8 cells was found to be the lowest for colchicine at 90 nM. Interestingly, microtubule polymerization inhibitors (colchicine and vinblastine) and dynein inhibitors (ciliobrevin D and dynarrestin) impaired ZFTA-RELA’s nuclear localization and NF-κB activity in 6E8 cells. These findings indicate that microtubule polymerization and dynein play pivotal roles in activating the NF-κB pathway in ST-EPN-ZFTA by promoting the nuclear localization of ZFTA-RELA. Consequently, inhibition of microtubule polymerization may be a therapeutic strategy for ST-EPN-ZFTA.

## INTRODUCTION

Ependymomas are rare primary brain and spinal tumors of the ependymal multiciliated epithelial cells that line the walls of the ventricles^1–3^. Based on their location, these tumors are classified into three types: supratentorial, posterior fossa, and spinal. A subclass of supratentorial ependymomas, termed ST-EPN-ZFTA, expresses multiple isoforms of a fusion protein consisting of a nuclear localizing protein with zinc finger domains, zinc finger translocation associated (ZFTA), and *v-rel* reticuloendotheliosis viral oncogene homolog A (RELA), an effector transcription factor of the nuclear factor-kappa B (NF-κB) pathway (hereafter referred to as ZFTA-RELA)^4–6^. To date, there has been no effective chemotherapy for ST-EPN-ZFTA.

NF-κB is an intracellular signaling pathway that transduces extracellular stimuli, such as stress and inflammatory cytokines, into cells and is involved in various processes, including tumorigenesis^7, 8^. In the absence of extracellular stimuli, RELA binds to IκB and localizes to the cytoplasm. Upon stimulation, IκB is phosphorylated and then degraded, and RELA translocates to the nucleus, where it regulates the expression of its target genes. Even in the absence of stimulation, ZFTA-RELA constitutively localizes to the nucleus and causes excessive activation of NF-κB signaling through the regulation of gene expression, ultimately leading to ST-EPN-ZFTA^4–6, 9–13^. Given the anticipated therapeutic potential of inhibitors of NF-κB activation by ZFTA-RELA for ST-EPN-ZFTA^14–16^, the development of high-throughput screening (HTS) systems and the search for lead compounds are of great importance.

It is anticipated that elucidating the nuclear transportation mechanism of ZFTA-RELA will facilitate an understanding of the pathogenic mechanism of ST-EPN-ZFTA. Given that ZFTA-RELA lacks only the initial three amino acids present in RELA and conserves all the functional domains of RELA^4–6^, it is speculated that the nuclear translocation machinery of RELA is also involved in that of ZFTA-RELA. Nevertheless, the precise mechanism by which ZFTA-RELA translocates to the nucleus remains unclear, and drugs that inhibit ZFTA-RELA’s nuclear migration have yet to be identified.

In this study, we develop an HTS system for compounds that inhibit NF-κB activation using ZFTA-RELA. Through the screening of 9600 compounds, we identify a derivative of colchicine as an NF-κB inhibitor. The results of this study suggest that pharmacological inhibition of microtubule polymerization and motor protein dynein, which transports its cargo along microtubules, impairs the nuclear localization of ZFTA-RELA and NF-κB activation. This study should contribute to the development of a novel treatment for ST-EPN-ZFTA by developing an HTS system and proposing novel therapeutic strategies.

## MATERIALS AND METHODS

### Cell culture

The NF-κB responsive luciferase reporter cell line 6E8, which expresses the most common isoform of ZFTA-RELA (ZFTA-RELA^FUS1^)^6^, has previously been established^14^. The HeLa/CMV-Luc cell line was established by Dr. T. Murakami at Saitama Medical University and purchased from the Japanese Collection of Research Bioresources (JCRB) at the National Institute of Biomedical Innovation, Health, and Nutrition (JCRB1679)^14^. HeLa cells were purchased from the American Type Culture Collection. These cell lines were cultured in Dulbecco’s modified Eagle medium (nacalai tesque) supplemented with 4.5 g/L glucose, 110 mg/L sodium pyruvate, 10% (v/v) fetal bovine serum (Biowest), and antibiotic-antimycotic solution (nacalai tesque), hereafter referred to as the culture media, at 37°C in 5% COL conditions^17, 18^. The expression of ZFTA-RELA in 6E8 cells was induced by the addition of 100 or 300 ng/mL doxycycline (Clontech) to the culture medium.

### Compounds

The DDI core library was provided by the Drug Discovery Initiative (DDI) of the University of Tokyo (https://www.ddi.f.u-tokyo.ac.jp/en/). The chemical compounds used in this study are listed in Supplementary Table 1.

### Relative viable cell counts and luciferase assays

The relative viable cell numbers were quantified using Cell Count Reagent SF (nacalai tesque) in accordance with the manufacturer’s instructions, using a TriStar LB 941 instrument (Berthold). For the luciferase assay, the compounds dissolved in dimethyl sulfoxide (DMSO) were added to 384-well flat-bottom white plates (Greiner Bio-One), and then 25 μL of 6E8, HeLa, or HeLa/CMV-Luc cell suspension adjusted to 250 cells/μL was added using the VIAFLO ASSIST system (Integra Biosciences). After overnight culture, luciferase assay was performed using TriStar LB 941 (Berthold) and the following reaction buffer: 100 mM Tris-HCl (pH 7.0, nacalai tesque), 5 mM MgCl_2_ (FUJIFILM), 250 μM coenzyme A trilithium salt (Oriental Yeast), 150 μM adenosine triphosphate (nacalai tesque), 150 μg/mL *D*-luciferin potassium salt (Cayman Chemical), 0.5 mM dithiothreitol (nacalai tesque), 50 μM ethylenediaminetetraacetic acid (pH8.0; Dojindo), and 0.1% Triton X-100 (Sigma-Aldrich). To calculate the inhibitory activities of the candidate compounds on the NF-κB responsive luciferase activity, the luciferase activity in 6E8 cells cultured in the presence or absence of doxycycline was defined as 0% or 100% inhibition, respectively. For the counter assay, the luciferase activity in HeLa/CMV-Luc or HeLa cells was similarly defined as 0% or 100% inhibition, respectively. The Z’-factor and S/N ratio were calculated according to a previous study^19^. IC_50_ values were calculated using the following formula:

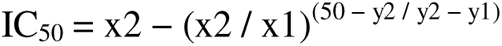

x1: sample concentration less than IC_50_,

x2: sample concentration higher than IC_50_,

y1: % inhibition at x1,

y2: % inhibition at x2.

### Quantitative PCR (qPCR)

cDNAs were prepared using NucleoSpin RNA (Macherey-Nagel) and ReverTra Ace (TOYOBO) kits, following the manufacturer’s instructions. Quantitative polymerase chain reaction (qPCR) was performed as reported previously^14, 15, 20^ using TB Green Premix Ex Taq II Tli RNaseH Plus (TaKaRa) on a StepOnePlus (Thermo Fisher Scientific). The primers utilized for qPCR are presented in Supplementary Table 2. *TATA binding protein* was used as an internal control. To calculate the inhibitory activities of the candidate compounds on gene expression, the mRNA expression levels in 6E8 cells cultured in the presence or absence of doxycycline were defined as 0% or 100% inhibition, respectively.

### Immunocytochemistry

The 6E8 cells were seeded on coverslips (Iwanami) coated with poly-*L*-lysine (nacalai tesque) and cultured overnight in the culture media supplemented with doxycycline and indicated compounds in 0.3% DMSO. The cells were then washed with Dulbecco’s phosphate-buffered saline (PBS) and fixed with 4% paraformaldehyde (PFA) dissolved in 100 mM phosphate buffer (PB, pH 7.4) at room temperature for 10 min. The fixed cells were washed with PBS and incubated in 10% normal goat serum (EMD Millipore) and 0.2% Triton X-100 (Sigma-Aldrich) in PBS (hereafter referred to as the blocking buffer). The cells were then incubated with mouse anti-FLAG antibody (Merck, clone M2) diluted in the blocking buffer at room temperature for 1 h, washed with PBS, incubated with donkey anti-mouse immunoglobulin G (IgG) conjugated with Alexa Fluor 488 (Abcam) and nuclear staining reagent [4’,6-diamidino-2-phenylindole (DAPI, Dojindo)] in PBS at room temperature for 1 h, washed with PBS, and mounted with Aqua-Poly/Mount (Polysciences). Confocal microscopy images were acquired using a Nikon AX. Nuclear localization ratios were calculated by dividing the degree of ZFTA-RELA-Flag immunoreactivity observed in the nucleus by that observed in the whole cell.

### Statistics

The means of three or more groups with one variable were compared with a one-way analysis of variance with Tukey’s post hoc test using Prism 9 (GraphPad). A *p*-value of less than 0.05 was considered statistically significant.

## RESULTS

### Establishment of an HTS system to identify compounds that inhibit NF-κB activity induced by ZFTA-RELA

Previously, we established the HEK293-derived cell line 6E8, which exhibits Flag-tagged ZFTA-RELA (ZFTA-RELA-Flag) expression and NF-κB responsive luciferase activity in a doxycycline dose-dependent manner^14^. Using this cell line, we identified terrein from *Aspergillus lentulus*, *epi*-aszonalenin B from *A. novofumigatus*, and aszonapyrone A from *Neosartorya spinosa* as NF-κB signaling inhibitors^14, 15^. To establish an HTS system to identify compounds that inhibit NF-κB signaling induced by ZFTA-RELA, we employed 384-well plates and automated dispensing machines (Fig. 1A and B). The Z’-factor and signal-to-background (S/B) ratio are frequently employed to assess the quality of an assay system^19^. A Z’-factor exceeding 0.5 is indicative of a robust and effective screening system. We calculated the S/B ratio and Z’-factor using the NF-κB-responsive luciferase activities from 6E8 cells cultured in medium without doxycycline as “backgrounds” and those from 6E8 cells cultured in medium with 100 ng/mL doxycycline as “signals.” The S/B ratio and Z’-factor of our assay were 219.14 ± 22.35 and 0.81 ± 0.05, respectively (mean ± standard deviation [SD], n = 30 plates; Fig. 1C and D). These results indicate that this assay is adaptable to HTS.

**Figure 1.**
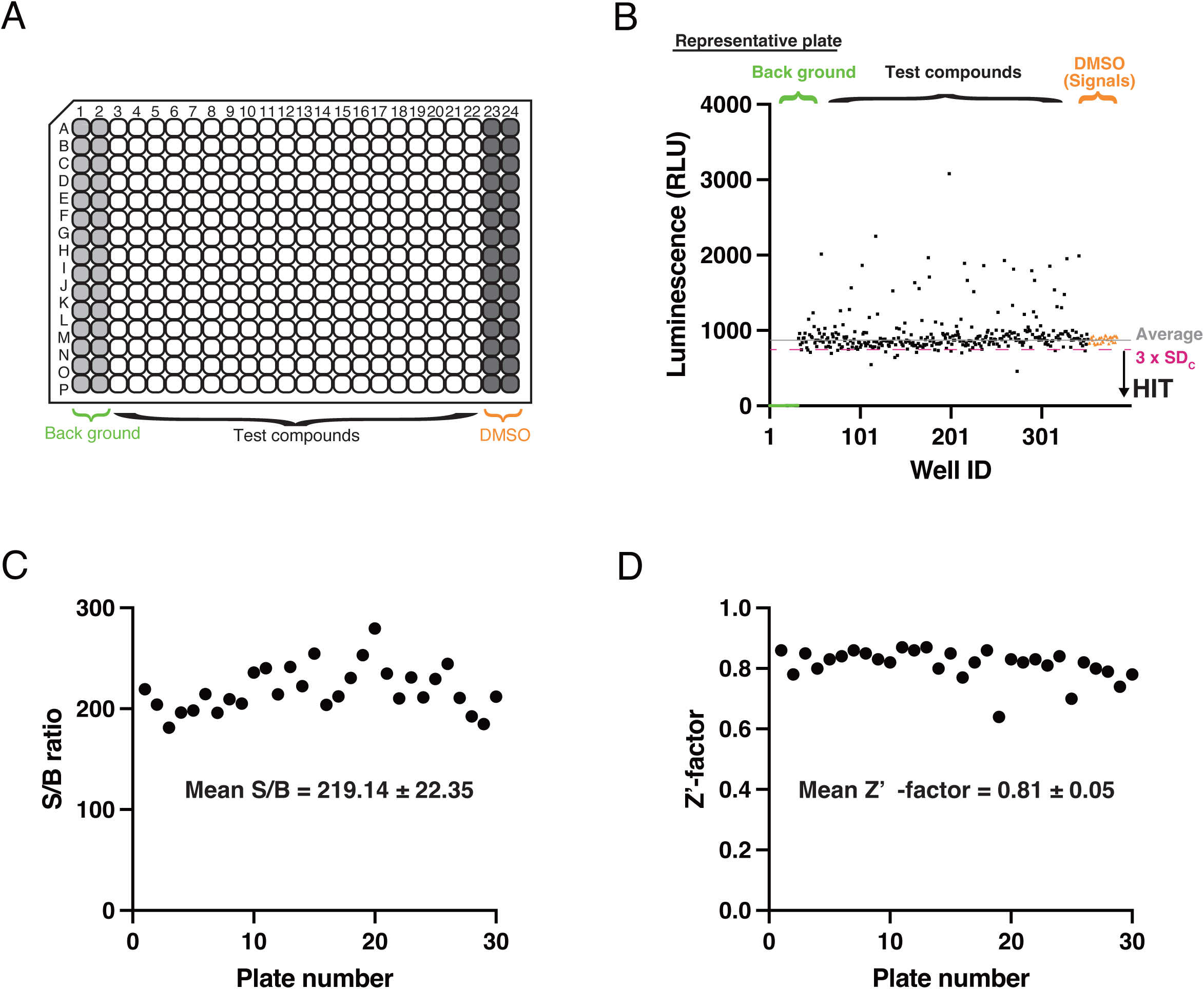
Establishment of an HTS system to identify compounds that inhibit NF-κB signaling induced by ZFTA-RELA. (A) Example of a 384-well plate with test compounds in the middle 352 wells and controls in the wells of the first and last two columns. (B) Representative plate analysis of the 384-well HTS. The graph shows the relative luminescent intensity of individual wells in a 384-well format. The samples with an intensity less than Mean_C_ − 3 × SD_C_ (mean and SD of DMSO controls, respectively) were identified as “HIT” in the primary screening. (C) Graph showing the signal-to-background ratio for each plate. (D) Graph showing the Z’-factor for each plate.

### Identification of a colchicine derivative as an inhibitor of NF-κB signaling induced by ZFTA-RELA

Using the HTS system, the DDI core library, consisting of 9600 chemicals selected for structural diversity, was screened. In the initial screening phase, the inhibitory activity of NF-κB was examined in a single well per compound at a concentration of 10 µM (Fig. 2A). The mean and SD of NF-κB-responsive luciferase activity were obtained from control wells with doxycycline, but no compounds were calculated for each plate (Mean_C_ and SD_C_, respectively). In the initial screening, 540 of the 9600 compounds demonstrated the capacity to inhibit NF-κB-responsive luciferase activity below the Mean_C_ – 3 × SD_C_ (shown as “HIT” in Fig. 1B). These 540 compounds were then subjected to a low-concentration study with a concentration of 5 µM per compound (Fig. 2A). At this concentration, 39 of the 540 compounds exhibited more than 50% inhibition of NF-κB-responsive luciferase activity upon the addition of doxycycline to 6E8 cells (shown as “HIT” in Fig. 2B), and these compounds were subjected to further tests.

**Figure 2.**
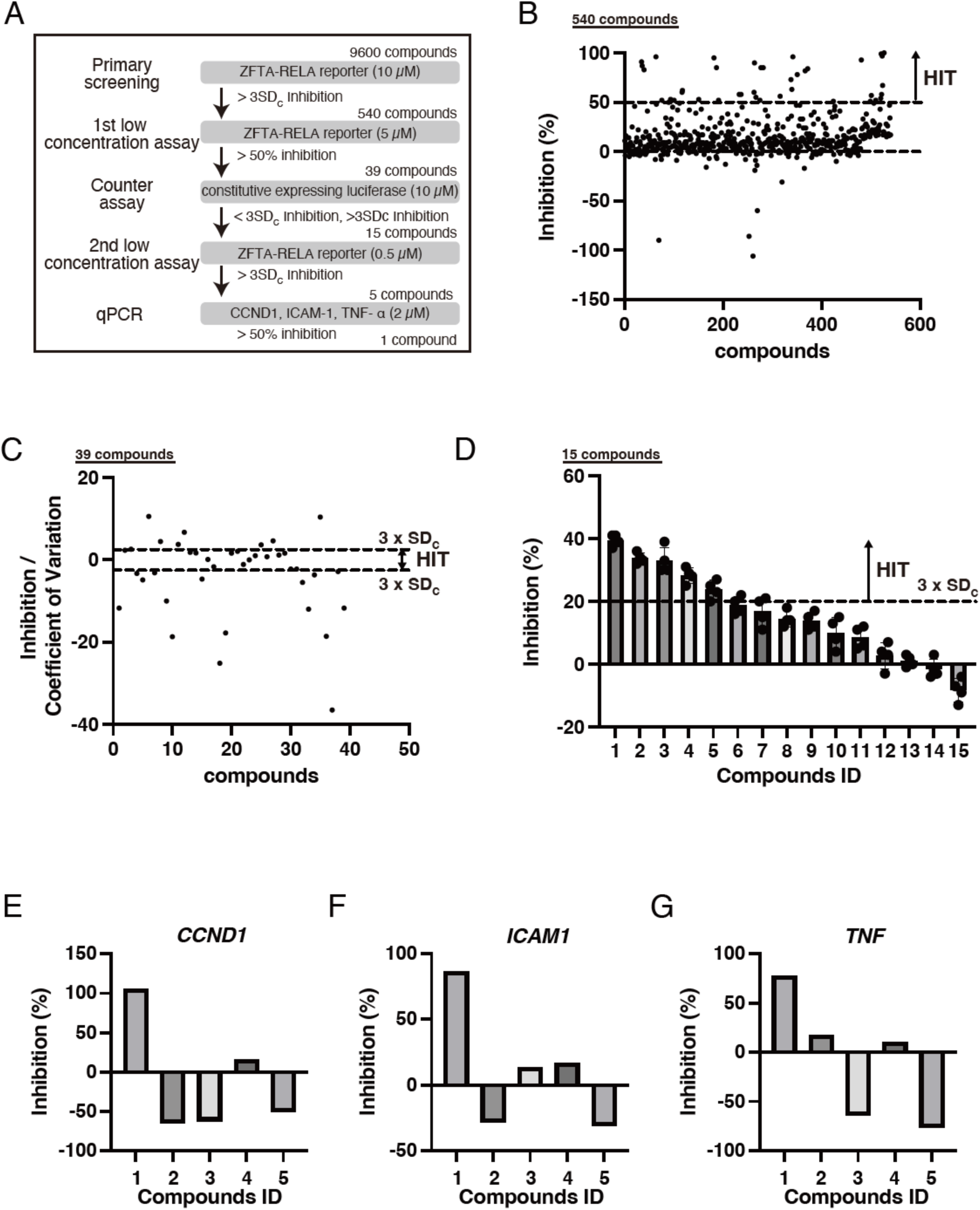
Identification of a colchicine derivative as an inhibitor of NF-κB signaling induced by ZFTA-RELA. (A) Scheme of the high-throughput screening strategy for the inhibitor of NF-κB signaling induced by ZFTA-RELA (B) Results of the first low-concentration assay, showing the effects of 540 compounds on NF-κB responsive luciferase activity induced by the expression of ZFTA-RELA in 6E8 cells. Each compound was added at a final concentration of 5 μM in DMSO. n = 4 each. The data shown are the mean inhibition rates of individual compounds. (C) The effects of 39 compounds on constitutively expressing luciferase activity in Hela/CMV-Luc cells (counter assay). Each compound was added at a final concentration of 10 μM in DMSO. n = 4 each. The data shown are the mean ± SD. The dots on the graph represent the average inhibition rates of individual samples. (D) The effects of compounds **1**–**15** on NF-κB responsive luciferase activity induced by expression of ZFTA-RELA in 6E8 cells. Each compound was added at a final concentration of 0.5 μM in DMSO. n = 4 each. The data shown are the mean ± SD. The dots on the graph represent the inhibition rates of individual samples. (E–G) Relative expression levels of *CCND1* (E), *ICAM1* (F), and *TNF* (G) mRNAs in 6E8 cells cultured overnight in the absence (-) or presence (+) of 100 ng/mL doxycycline. The mRNA level of the doxycycline control was set to 100%. Compounds **1**–**5** dissolved in DMSO were added to 6E8 cells cultured in the presence of doxycycline. n = 1 each.

The employment of 6E8 cells in the compound screening process may lead to the identification of compounds that exhibit a nonspecific inhibitory effect on luciferase reaction and/or cell viability^14, 15^. To exclude nonspecific inhibitory compounds from those obtained by the previous screening process, a counter assay at 10 µM was performed using HeLa/CMV-Luc cells that constitutively expressed luciferase under the cytomegalovirus (CMV) enhancer/promoter (Fig. 2A). The Mean_C_ and SD_C_ of constitutive luciferase activity obtained from HeLa/CMV-Luc cells were calculated. For 15 of the 39 compounds, the constitutive luciferase activity was within the Mean_C_ ± 3 × SD_C_ (Fig. 2A and C). These compounds were deemed to exhibit no nonspecific activation or inhibition of constitutive luciferase activity and were therefore submitted to the next test (shown as “HIT” in Fig. 2C and named as compounds **1**–**15**). A second low-dose study was performed using a sample with a concentration of 0.5 µM to select compounds that could be effective at low doses (Fig. 2A). Five of the 15 compounds showed more than 20% luciferase inhibitory activity (corresponding to the NF-κB-responsive luciferase activity being below Mean_C_ – 3 × SD_C_) under these low-concentration conditions (compounds **1**–**5**, shown as “HIT” in Fig. 2D).

The expression of three endogenous NF-κB responsive genes, namely *CCND1*, *ICAM1*, and *TNF*, increases in a doxycycline-dependent manner in 6E8 cells^14, 15^. The effects of compounds **1**–**5** on the expression of these three endogenous NF-κB responsive genes were examined using qPCR. Subsequently, the inhibitory activities of these five compounds on the expression of the three genes were calculated (Fig. 2E–G). Of the compounds tested, compound **1**, a derivative of colchicine that inhibits microtubule polymerization^21^, demonstrated the greatest inhibitory activity on the expression of the three endogenous NF-κB response genes (Fig. 2E–G). Consequently, the subsequent analysis focused on colchicine and its derivatives.

### Colchicine and its derivatives inhibit the NF-κB-responsive reporter activity induced by ZFTA-RELA

Following the aforementioned screening, which yielded colchicine derivative compound **1**, we prepared colchicine and six derivatives of colchicine (compounds **1** and **16**–**20**) and conducted a comparative analysis of their NF-κB inhibitory activity using 6E8 cells (Fig. 3A and B). Although colchicine exhibited the lowest IC_50_, with a value of 0.090 µM, C7-derivatives of colchicine, including compounds **1**, **16** (*N*-deacetyl-*N*-methylcolchicine), and **17** (speciosine), also demonstrated inhibitory activity within the submicromolar range (0.22–0.59 μM). In contrast, the inhibitory activities against ZFTA-RELA-dependent luciferase activity were significantly diminished (3.0–5.2 μM) by the substitution of the C10-methoxy group with alkylamino or iminium moieties, as exemplified by compounds **18**–**20**. In the following study, we focus on colchicine because it has the highest NF-κB inhibitory activity (lowest IC_50_) and is a pharmaceutical product.

**Figure 3.**
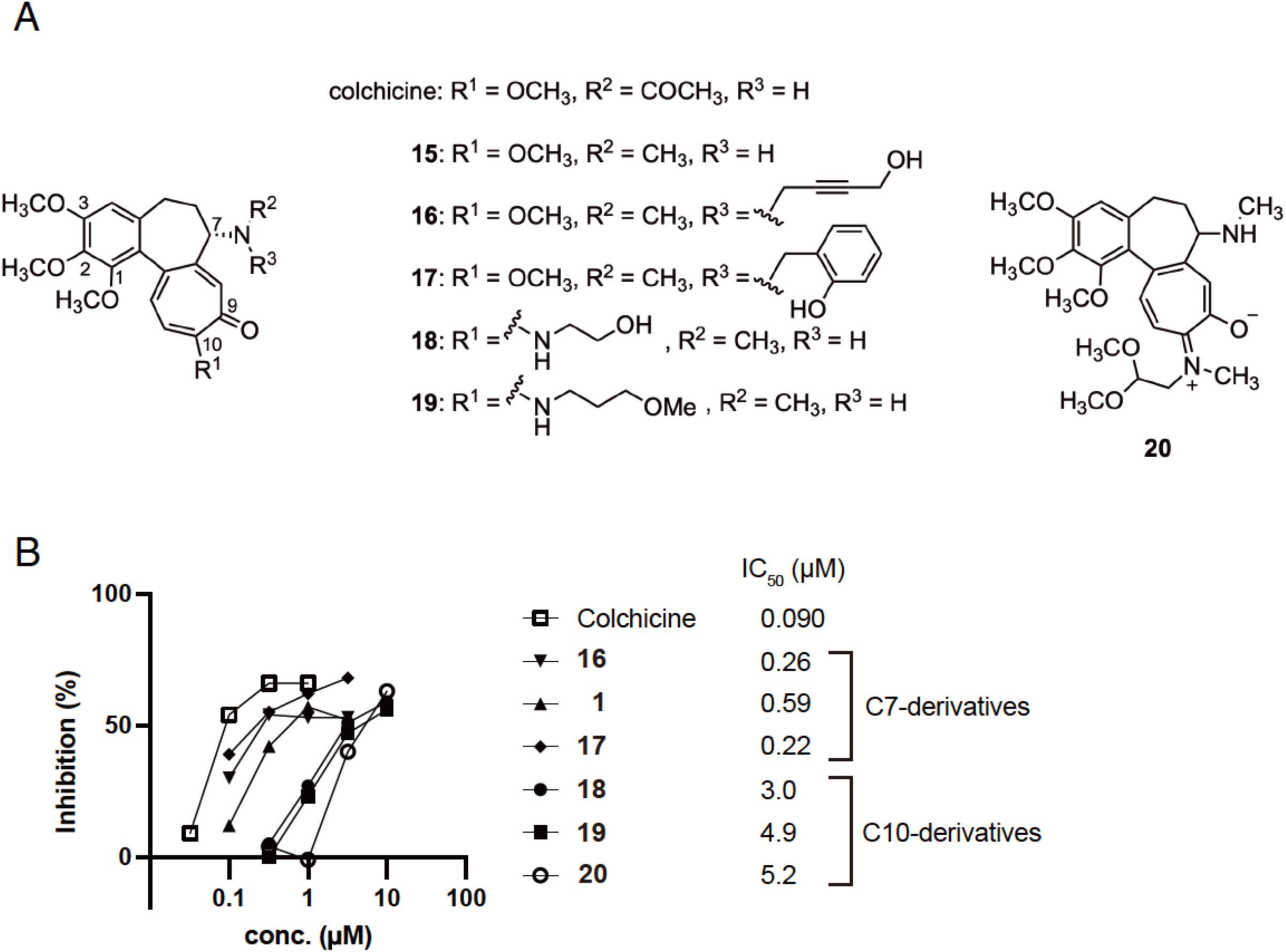
Comparison of the inhibitory activity of colchicine derivatives on NF-κB-responsive luciferase activity induced by ZFTA-RELA. (A) Structures of colchicine and its derivatives (compounds **1** and **16**–**20**) used in the present study. (B) Titration assays of colchicine and colchicine derivatives in the inhibition of ZFTA-RELA-induced NF-κB responsive luciferase activities.

### Colchicine inhibition of endogenous NF-κB-responsive gene expression induced by ZFTA-RELA

To confirm the screening results, we examined the effect of colchicine on the upregulation of endogenous NF-κB response gene expression by expression of ZFTA-RELA using the 6E8 cells. As previously reported^14, 15^, the addition of doxycycline to the 6E8 cells resulted in increased expression of *CCND1*, *ICAM1*, *TNF*, and *L1CAM* (Fig. 4A–D). Colchicine inhibited the upregulation of these gene expressions (Fig. 4A–D). An endogenous NF-κB responsive gene expression study revealed that overexpression of ZFTA-RELA, but not RELA, results in the upregulation of *ZFTA* expression^13, 20^. Notably, colchicine inhibited the upregulation of *ZFTA* expression induced by ZFTA-RELA (Fig. 4E). These results suggest that colchicine inhibits the processes that lead to transcriptional activation by ZFTA-RELA, including its nuclear transportation.

**Figure 4.**
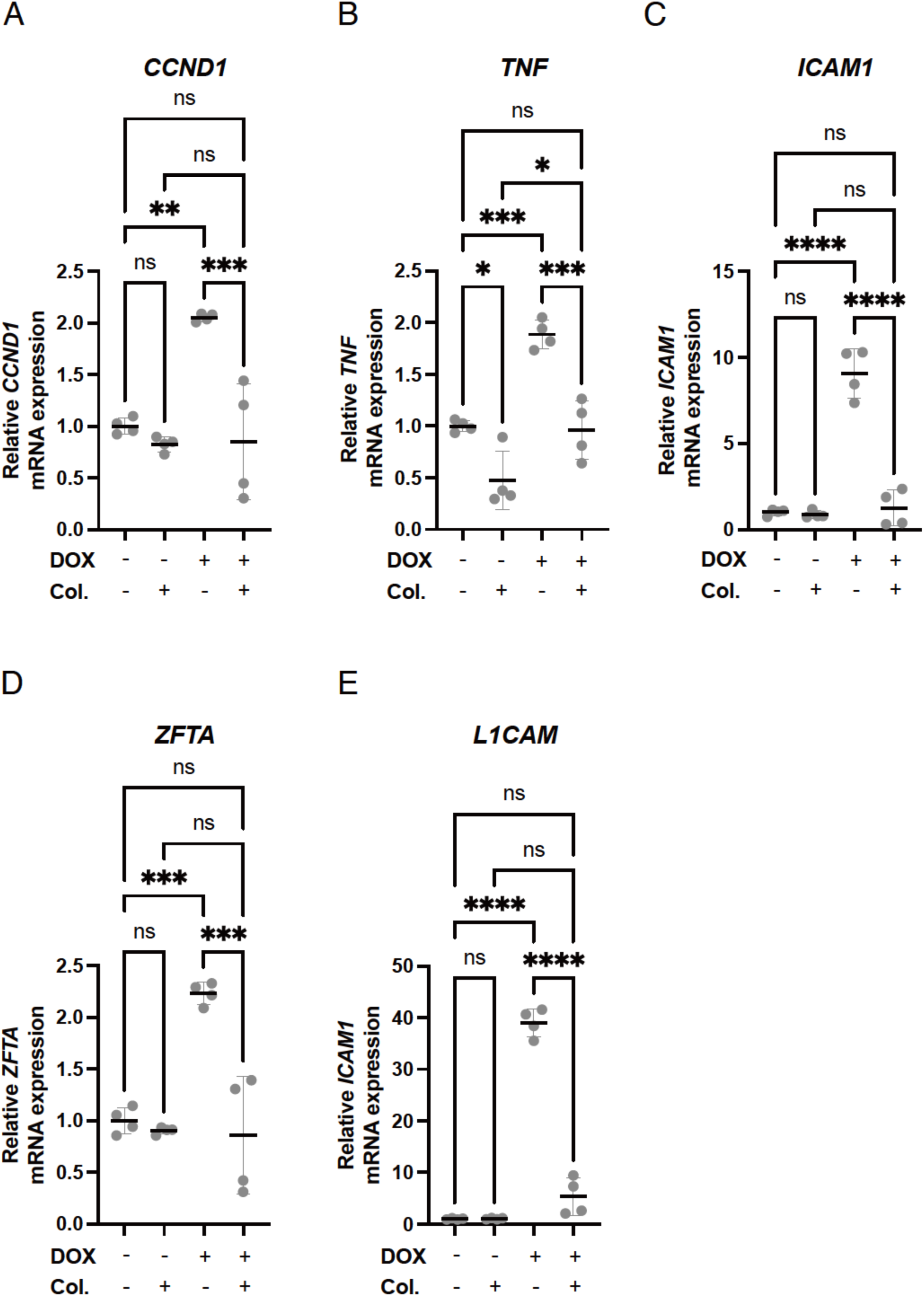
Effect of colchicine on the expression of endogenous NF-κB responsive genes and *ZFTA*. (A–E) Relative expression levels of *CCND1* (A), *ICAM1* (B), *TNF* (C), *L1CAM* (D), and *ZFTA* (E) mRNAs in 6E8 cells cultured overnight in the absence (-) or presence (+) of 300 ng/mL doxycycline (DOX) and 0.3 µM colchicine (Col.). The mRNA level of the doxycycline control was set to 1. The data shown are the mean ± SD. Each point on the graph represents the relative mRNA expression level of an individual sample. n = 4 each. ns, not significant; *p < 0.05; **p ≤ 0.005; ***p ≤ 0.0005; ****p < 0.0001.

### Involvement of microtubules and dynein in the NF-κB activation induced by ZFTA-RELA

ZFTA-RELA is constitutively localized to the nucleus, even in the absence of stimuli that activate the NF-κB pathway^6, 10, 11, 13^. In the 6E8 cells added with DMSO and doxycycline (DMSO control), ZFTA-RELA-Flag was localized to the nucleus (Fig. 5A). In contrast, the nuclear localization of ZFTA-RELA-Flag was inhibited in the 6E8 cells to which 0.10 µM colchicine was added (Fig. 5A). The inhibition of nuclear localization of ZFTA-RELA-Flag in the 6E8 cells by colchicine was confirmed by calculating the nuclear localization ratio (nuclear ZFTA-RELA-Flag immunoreactivity / that of the whole cell) (Fig. 5B). These findings suggest that colchicine inhibits the nuclear localization of ZFTA-RELA, thereby blocking the expression of ZFTA-RELA-responsive genes.

**Figure 5.**
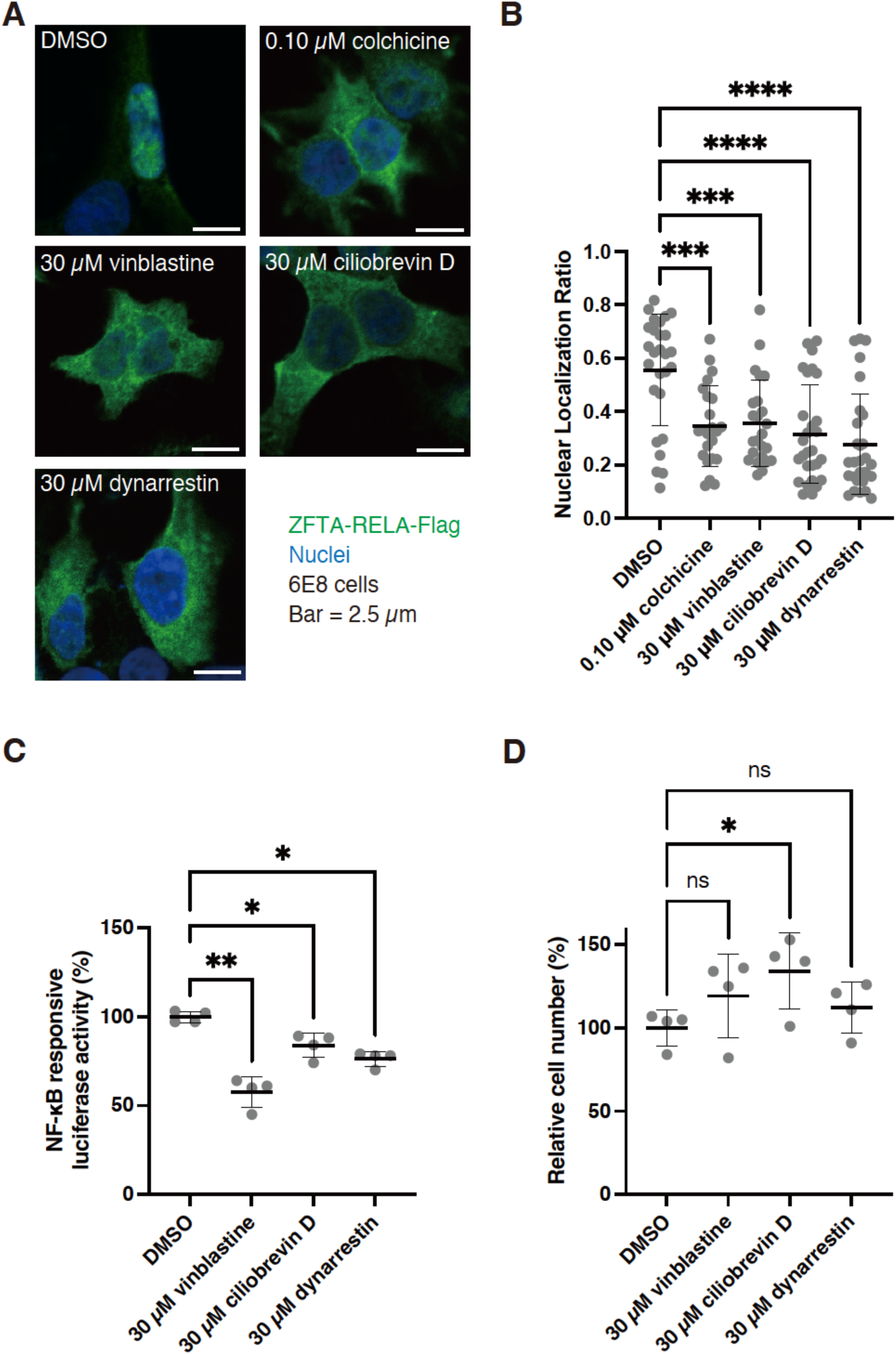
The vinblastine, ciliobrevin D, and dynarrestin treatments resulted in reductions in NF-κB-responsive luciferase activity and the nuclear localization ratio of ZFTA-RELA in 6E8 cells. (A, B) Representative confocal images (A) and quantification of the nuclear localization of ZFTA-RELA-Flag (B) in 6E8 cells. ZFTA-RELA-Flag (green), nuclei (blue). DMSO served as the solvent control. The scale bar represents a length of 2.5 µm. The data presented are the mean ± SD. Each data point on the graph represents the nuclear localization ratio of an individual cell. The number of samples in each group was as follows: DMSO, n = 27; colchicine, n = 23; vinblastine, n = 24; ciliobrevin D, n = 28; and dynarrestin, n = 28. ***p < 0.001; ****p < 0.0001. (C, D) NF-κB-responsive luciferase activity (C) and the relative viable cell number (D) in the 6E8 cells. DMSO served as the solvent control. The mean reporter activity (C) and cell number (D) in DMSO control cells were set to 100%. The data presented are the mean ± SD. Each data point on the graph represents the relative reporter activity (C) and cell number (D) of an individual sample. The number of samples (n) was four for each data point. ns, not significant; *p < 0.05; **p < 0.01.

It has been reported that the destabilization of microtubules inhibits the activation of the NF-κB pathway and the nuclear translocation of RELA induced by TNF-α stimulation^22^. Dynein is a motor protein that moves along microtubules, facilitating the nuclear transport of RELA^23^. To ascertain whether microtubule polymerization and dynein are also involved in the activation of the NF-κB pathway by the expression of ZFTA-RELA and the nuclear localization of ZFTA-RELA, we employed another microtubule polymerization inhibitor, vinblastine, and two inhibitors of dynein, ciliobrevin D and dynarrestin, in the 6E8 cells^24–26^. The nuclear localization of ZFTA-RELA-Flag was inhibited in the 6E8 cells to which 30 µM vinblastine, 30 µM ciliobrevin D, or 30 µM dynarrestin was added (Fig. 5A and B). Furthermore, the administration of vinblastine, ciliobrevin D, and dynarrestin demonstrated an inhibitory effect on an increase in NF-κB-responsive luciferase activity by the addition of doxycycline to the culture media of the 6E8 cells (Fig. 5C). On the other hand, these inhibitors did not reduce the number of viable cells in the 6E8 cells (Fig. 5D). These results suggest that microtubule polymerization plays a pivotal role in the nuclear localization of ZFTA-RELA, which in turn activates the NF-κB pathway.

## DISCUSSION

In this study, we developed an HTS system for the exploration of potential therapeutic compounds for ST-EPN-ZFTA, for which no effective chemotherapy has yet been established. The HTS and subsequent analysis demonstrated that the microtubule polymerization inhibitor colchicine inhibited the NF-κB activity induced by ZFTA-RELA. Furthermore, our findings demonstrate the importance of microtubule polymerization and dynein in the nuclear migration of ZFTA-RELA, which results in the hyperactivation of the NF-κB pathway. Through the establishment of the HTS system, this study not only proposes a new treatment strategy for ST-EPN-ZFTA but also elucidates a part of the molecular mechanisms underlying the pathogenesis of ST-EPN-ZFTA.

In this study, we developed an HTS system to identify inhibitors of ZFTA-RELA-induced NF-κB-responsive luciferase reporter activity and to identify colchicine derivatives. The luciferase assay is a suitable method for HTS because it is simple and quantitative. The S/B ratio and Z’-factor of our HTS indicate that it is a reliable screening system. A counter assay was conducted using HeLa/CMV-Luc cells to identify compounds that reduce NF-κB-responsive luciferase activity in 6E8 cells due to cytotoxicity and/or nonspecific inhibition of the luciferase assay. Since the CCK-8 assay, in which a dye is produced by NADH-mediated redox reactions in living cells, can also exclude compounds with cytotoxic activity, this assay was performed after the HeLa/CMV-Luc counter-assay. However, no additional compounds were excluded (data not shown). Thus, it would be sufficient to use a method that is more convenient for the researcher.

In this study, five compounds were identified in HTS to inhibit NF-κB-responsive luciferase activity in 6E8 cells. However, three out of five did not inhibit the expression of all three endogenous NF-κB-responsive genes tested (Fig. 2E–G). This discrepancy may be attributed to differences in sensitivity between the luciferase assay and qPCR. These compounds may be active in inhibiting the expression of endogenous NF-κB-responsive genes when the degree of NF-κB activation is lower. An alternative hypothesis is that the artificial NF-κB responsive elements in 6E8 cells do not fully reflect the complexity of NF-κB activity. In this case, subsequent screening would require a multifaceted investigation that includes consideration of the expression of endogenous responsive genes, as demonstrated in the present study.

Colchicine is primarily employed in the management of gout and familial Mediterranean fever due to its anti-inflammatory properties^21^. Nevertheless, it is not a commonly utilized treatment for cancer due to its toxicity^27–29^. While our findings indicate that colchicine inhibits NF-κB activity induced by ZFTA-RELA, the development of effective and less toxic colchicine derivatives is a crucial subsequent step. Consistent with previous reports^27, 30, 31^, the C7-derivatives exhibited a minimal decline in NF-κB inhibitory activity, while the C10-derivatives demonstrated a pronounced reduction in inhibitory activity (Fig. 3). Considering that the C10-methoxy group of colchicine is located on the binding side with tubulin^32, 33^, it is postulated that C10-derivatives may exhibit diminished efficiency in binding to tubulin due to steric hindrance. A more promising route for the synthesis of colchicine derivatives would be the modification of C7, given that C7 is on the opposite side to the tubulin binding side^32, 33^ and that the NF-κB inhibitory activity was not significantly reduced in the C7-derivatives. Furthermore, the nitrogen at C7 has been demonstrated to be indispensable for the recognition of colchicine derivatives by P-glycoprotein, which facilitates the expulsion of drugs from the cell^34^.

Given that all the functional domains of RELA are conserved in ZFTA-RELA^6^, it is anticipated that ZFTA-RELA is transported to the nucleus, at least in part, by the same mechanism as RELA. Here, we showed that inhibition of microtubule polymerization and dynein disrupts the nuclear localization of ZFTA-RELA, in accordance with previous reports on RELA’s nuclear transportation. First, colchicine inhibits TNF-α-and cleaved pyrine-stimulated nuclear transportation of RELA and activation of the NF-κB pathway, while the microtubule-stabilizing agent taxol activates the NF-κB pathway^35, 36^. Second, the knockdown of dynein, a motor protein that transports cargo along microtubules, inhibits the nuclear translocation of RELA^23^. Given that RELA binds to dynein via importin and is transported from the axon along microtubules to the cell body in hippocampal neurons^37^, ZFTA-RELA may be similarly transported to the nucleus by dynein by binding to importin.

ZFTA-RELA would have been transported to the nucleus by additional mechanisms that do not involve microtubule polymerization or dynein because pharmacological inhibition of microtubule polymerization or dynein did not completely inhibit nuclear translocation of ZFTA-RELA or NF-κB-responsive luciferase activity (Fig. 3B). This idea is supported by previous reports that the fusion of a partial sequence of ZFTA to GFP localizes it to the nucleus and that deletion of the zinc finger domain from the ZFTA portion of ZFTA-RELA inhibits its nuclear localization^10, 11^. Further studies are needed to address the molecular mechanisms underlying the nuclear transportation of ZFTA-RELA, which leads to the pathogenesis of ST-EPN-ZFTA.

Through the development of an HTS system, this study provides a novel strategy for the treatment of ST-EPN-ZFTA as well as a comprehensive understanding of the underlying molecular mechanisms that contribute to its pathogenesis. This study is expected to inform future research needed to overcome difficulties in the treatment of ST-EPN-ZFTA.

## Abbreviations

ST-EPN-ZFTA: supratentorial ependymoma, ZFTA-fusion positive
ZFTA: zinc finger translocation associated
ZFTA-RELA: fusion protein of ZFTA and RELA

## Supporting Information

Supplementary Table 1. Chemical compounds used in the present study Supplementary Table 2. Primers used in the present study

## Acknowledgments

The authors would like to express their gratitude to Dr. T. Murakami of Saitama Medical University and the JCRB for providing the HeLa/CMV-Luc cell line, and K. Egashira, M. Ichihara, M. Kikuchihara, R. Komuro, and H. Uga for their technical assistance.

