## Supplemental Tables for "Inhibition of microtubule polymerization and dynein impairs the nuclear localization of the ependymoma-associated ZFTA-RELA fusion protein and NF-κB activation"

Supplementary Table 1. Chemical compounds used in the present study

| Compounds | Supplier | Supplier ID | SMILES |
| --- | --- | --- | --- |
| Ciliobrevine D | Selleck | S9743 |  |
| Colchicine | FUJIFILM | C5645 |  |
| Dynarrestin | Cayman | 35527 |  |
| Vinblastine | Angene | AG019FX5 |  |
| **1** | ChemDiv | N051-0027 | COC1=CC=C2C(=CC1=O)[C@H](CCc3cc(OC)c(OC)c(OC)c23)N(C)CC#CCO |
| **2** | Enamine | T5325727 | CC(Nc1ccccc1CO)C(=O)Nc2cccc(Cl)c2Cl |
| **3** | Vitas-M | STK047386 | COc1ccc2c(c1)[nH]c3c(C)nccc23 |
| **4** | Vitas-M | TULIP022047 | CCc1c(c2ccccc2)n(CCO)c3nc4N(C)C(=O)N(C)C(=O)c4n13 |
| **5** | LifeChemicals | F0578-0280 | CCN(c1ccc(OC)cc1)c2ncnc3[nH]cnc23 |
| **6** | Enamine | T5446166 | CS(=O)(=O)c1ccc(cc1[N+](=O)[O-])C(=O)Nc2nc3ccccc3[nH]2 |
| **7** | Pharmeks | P2001S-273238 | COc1ccc(cc1OC)C(=O)N(CC(C)C)CC2=Cc3ccc(C)cc3NC2=O |
| **8** | Pharmeks | P2000N-27762 | C[C@H]1CCC[C@]2(C)C[C@H]3OC(=O)C(CNC(C)(CO)CO)[C@H]3C=C12 |
| **9** | Pharmeks | P2001S-230553 | COc1ccc(cc1)c2cn[nH]c2c3ccc(O)cc3O |
| **10** | Enamine | T6120521 | CC(OC(=O)c1ccc(c(c1)[N+](=O)[O-])S(=O)(=O)C)C(=O)N2CCC(C)CC2 |
| **11** | ChemDiv | K072-0101 | Clc1cncc(Cl)c1N\N=C\c2ccncc2 |
| **12** | ChemDiv | G943-0146 | CN1C(=O)c2ccccc2Sc3cc(NC(=O)c4ccnn4C)ccc13 |
| **13** | Enamine | T5559448 | CCCCN1C(=O)NC(=O)c2c1nc(COC(=O)C(C(C)CC)c3ccccc3)n2CC |
| **14** | Vitas-M | STK289683 | CC(=O)NC1=NN(C(=O)C)C(C)(S1)c2ccc(Cl)s2 |
| **15** | Zelinsky Institute | UZI/2320824 | C(c1ccccc1)n2c3c(CCc4nonc34)c5ccccc25 |
| **16** | Alexis | 430-033 | CNC1CCc2cc(OC)c(OC)c(OC)c2C3=CC=C(OC)C(=O)C=C13 |
| **17** | ChemDiv | N051-0003 | COC1=CC=C2C(=CC1=O)[C@H](CCc3cc(OC)c(OC)c(OC)c23)N(C)Cc4ccccc4O |
| **18** | Pharmeks | P2000N-06164 | CN[C@H]1CCc2cc(OC)c(OC)c(OC)c2C3=CC=C(NCCO)C(=O)C=C13 |
| **19** | Pharmeks | P2000N-21042 | CN[C@H]1CCc2cc(OC)c(OC)c(OC)c2C3=CC=C(NCCCOC)C(=O)C=C13 |
| **20** | Pharmeks | P2000N-28889 | CNC1CCc2cc(OC)c(OC)c(OC)c2C3=C1C=C([O-])\C(=[N+](/C)\CC(OC)OC)\C=C3 |

Supplementary Table 2. Primers used in the present study

| Primer Name | Sequence | PrimerBank ID |
| --- | --- | --- |
| *CCND1* F | 5′-CAATGACCCCGCACGATTTC-3′ | 77628152c3 |
| *CCND1* R | 5′-CATGGAGGGCGGATTGGAA-3′ |  |
| *ICAM1* F | 5′-ATGCCCAGACATCTGTGTCC-3′ | 167466197c1 |
| *ICAM1* R | 5′-GGGGTCTCTATGCCCAACAA-3′ |  |
| *L1CAM* F | 5′-TGTCATCACGGAACAGTCTCC-3′ | 221316755c1 |
| *L1CAM* R | 5′-CTGGCAAAGCAGCGGTAGAT-3′ |  |
| *TBP* F | 5′-GAGCCAAGAGTGAAGAACAGTC-3′ | 285026518c3 |
| *TBP* R | 5′-GCTCCCCACCATATTCTGAATCT-3′ |  |
| *TNF* F | 5′-CCTCTCTCTAATCAGCCCTCTG-3′ | 25952110c1 |
| *TNF* R | 5′-GAGGACCTGGGAGTAGATGAG-3′ |  |
| *ZFTA* F | 5′-GGGTCTGGAGGAAGAGATGCC-3′ |  |
| *ZFTA* R | 5′-TGCCCTCTGCTTTCCCACTCC-3′ |  |
